# Pouch microbiome changes during lactation in the short-beaked echidna (*Tachyglossus aculeatus*)

**DOI:** 10.1101/2024.02.28.581309

**Authors:** Isabella Wilson, Tahlia Perry, Raphael Eisenhofer, Peggy Rismiller, Michelle Shaw, Frank Grutzner

## Abstract

Monotreme and marsupial early development is characterised by a short gestation, the birth of the young at an early stage of development, and a long lactation in the pouch or pseudo-pouch. The lack of a functional adaptive immune system in these altricial young raises questions about how they survive in a microbe-rich world. Previous studies on marsupial pouches have revealed changes to pouch microbe composition during lactation but no information is available in monotremes. We investigated changes in the echidna pseudo-pouch microbiome during different stages of the reproductive cycle and whether this differs between wild and zoo-managed animals. Swab samples were obtained from wild and captive echidna pseudo-pouches outside of breeding season, during courtship and breeding, and during lactation. 16S rRNA gene metabarcoding revealed that the pseudo-pouch microbiome undergoes dramatic changes during lactation, with a reduction in bacterial taxa that may be pathogenic. These changes were not observed in samples taken outside of breeding season or during courtship and mating. This showed that the echidna pseudo-pouch environment changes during lactation to accommodate young that lack a functional adaptive immune system. This study pioneers pouch microbiome research in monotremes, provides new biological information on echidna reproduction, and may also provide information about the effects of captive management to inform breeding programs in the future.

## Introduction

The reproductive microbiome, which includes vaginal, milk, and mammary microbiota, is increasingly being recognised for its contributions to infant health^[1]^. The first major exposure to bacteria occurs during birth when the infant comes into contact with maternal vaginal, faecal, and skin microbiota^[2]^. This results in colonisation of oral, skin, nasopharyngeal, and gastrointestinal habitats within the infant^[3]^. Subsequent exposure through lactation, including microbes from the milk as well as the skin surrounding the nipple, is crucial for the colonisation of the infant gut microbiome^[1]^. Milk provides antimicrobials that prevent the growth of pathogenic species, and in humans, also contains prebiotic oligosaccharides to support the growth of commensal microbes^[4,5]^.

Some of these maternally-derived microbes may play critical roles in early development, including brain development, motor control, immune function, metabolism, and homeostasis^[6,7]^. It is unsurprising, then, that the microbial environment in which neonates develop is of great importance to health and disease; delivery via caesarean section is associated with infant gut dysbiosis and an increased risk of obesity and immune-related disorders such as type 1 diabetes, asthma, and allergy^[8–12]^.

In monotremes and marsupials, the reproductive microbiome extends to the pouch – an environment critical for the development of their altricial young. Research into the marsupial pouch microbiome has revealed compositional changes after the birth of the joey^[13]^. These changes are thought to be the result of the secretion of anti-microbial agents in the skin and milk, including cathelicidin, lysozyme, dermcidin, and immunoglobulins^[14,15]^. This protective activity appears to modulate the pouch microbiota, potentially bringing about more favourable conditions for the young^[16]^. While similar secretions have been identified in monotremes^[17]^, the reproductive microbiome has not yet been studied.

Monotremes offer a unique perspective on reproduction and early development as the only egg-laying mammals. The embryo undergoes a short gestation resulting in an altricial hatchling which lacks a functioning adaptive immune system^[18,19]^. The majority of early development occurs during a long lactation (∼160-210 days in echidnas)^[20]^ within the pseudo-pouch, a temporary pouch formed from the contraction of the abdominal muscles. Monotremes lack nipples, instead nursing from the milk patch within the pseudo-pouch. Nothing is currently known about the monotreme pseudo-pouch microbiome.

This study investigates for the first time how seasonal changes and lactation affect the pseudo-pouch microbiota of wild and captive short-beaked echidnas. Wild and captive echidnas were swabbed during three different reproductive stages: outside of breeding season, within breeding season (June-September), and during lactation. This revealed extensive microbial changes within the pseudo-pouch after the onset of lactation.

## Results

The lack of a functioning adaptive immune system in monotreme hatchlings raises questions about the pseudo-pouch microbiome during lactation. We hypothesised that secretions during breeding season or lactation would affect microbes in the pseudo-pouch, similar to what has been observed in the marsupial pouch. We performed 16S rRNA gene sequencing on a total of 63 samples: pseudo-pouch (n = 22), cloacal (n = 17), oral (n = 20), and habitat (n = 4). Swab samples were taken from wild and captive echidnas during three stages of the reproductive cycle (Table 1).

**Table 1:** Overview of samples extracted within this study, sorted by sample type and species.

| Sample type | Captive | Wild | Total |
| --- | --- | --- | --- |
| Pouch (lactating) | 6 | - | 6 |
| Pouch (inside of breeding season) | - | 4 | 4 |
| Pouch (outside of breeding season) | 4 | 8 | 12 |
| Oral | 9 | 11 | 20 |
| Cloacal | 7 | 10 | 17 |
| Environment | 3 | 1 | 4 |
| **Total** | **29** | **38** | **63** |

### Decontamination of the dataset

As DNA contamination can confound microbiome studies, and the monotreme pseudo-pouch microbiome has not previously been studied, we first sought to validate the presence of microbes in the echidna pseudo-pouch. We used the decontam R package to identify putative contaminants within the dataset (Supp Figure S7; Supp Figure S8). This approach compares negative controls to biological samples in order to classify taxa as contaminants or non-contaminants. We captured contaminants introduced during the DNA extraction process using extraction blanks (i.e. samples to which no echidna swabs were added; n = 10) and during PCR using negative controls (i.e. PCR reactions to which no DNA was added; n = 10). Using a prevalence-based method with a threshold score of 0.5, 586 features were identified as contaminants out of a total of 12,209 features. These were removed from the dataset prior to further analyses.

### Lactation-associated changes in the pseudo-pouch microbiome of wild and captive short-beaked echidnas

We compared the diversity and taxonomic composition of pseudo-pouch microbiota samples from lactating and non-lactating echidnas. Microbial diversity was analysed using four different alpha diversity metrics: species richness (i.e. the number of species, measured using observed OTUs and Shannon’s index), species evenness (i.e. the distribution of species abundances, measured using Pielou’s evenness), and phylogenetic diversity (a measure of diversity which incorporates branch length, measured using Faith’s phylogenetic diversity).

Phylogenetic diversity was significantly reduced within the lactating pseudo-pouch (Figure 1a; p = 0.022) as was species richness as measured by observed OTUs (Figure 1b; p = 0.022). Species evenness and Shannon’s diversity were not significantly affected by lactation status (Supp Fig S1; p > 0.05). Analysis of beta diversity revealed that lactating pseudo-pouch samples were broadly dissimilar from their non-lactating counterparts, forming a distinct cluster across axes 1 and 2 during principal coordinate analysis (PCoA; Figure 1c). There was some cross-over with two of the lactating samples falling within the non-lactating cluster – however, the microbial composition of the pseudo-pouch was found to be significantly different between lactating and non-lactating samples (weighted UniFrac; p = 0.015).

**Figure 1:**
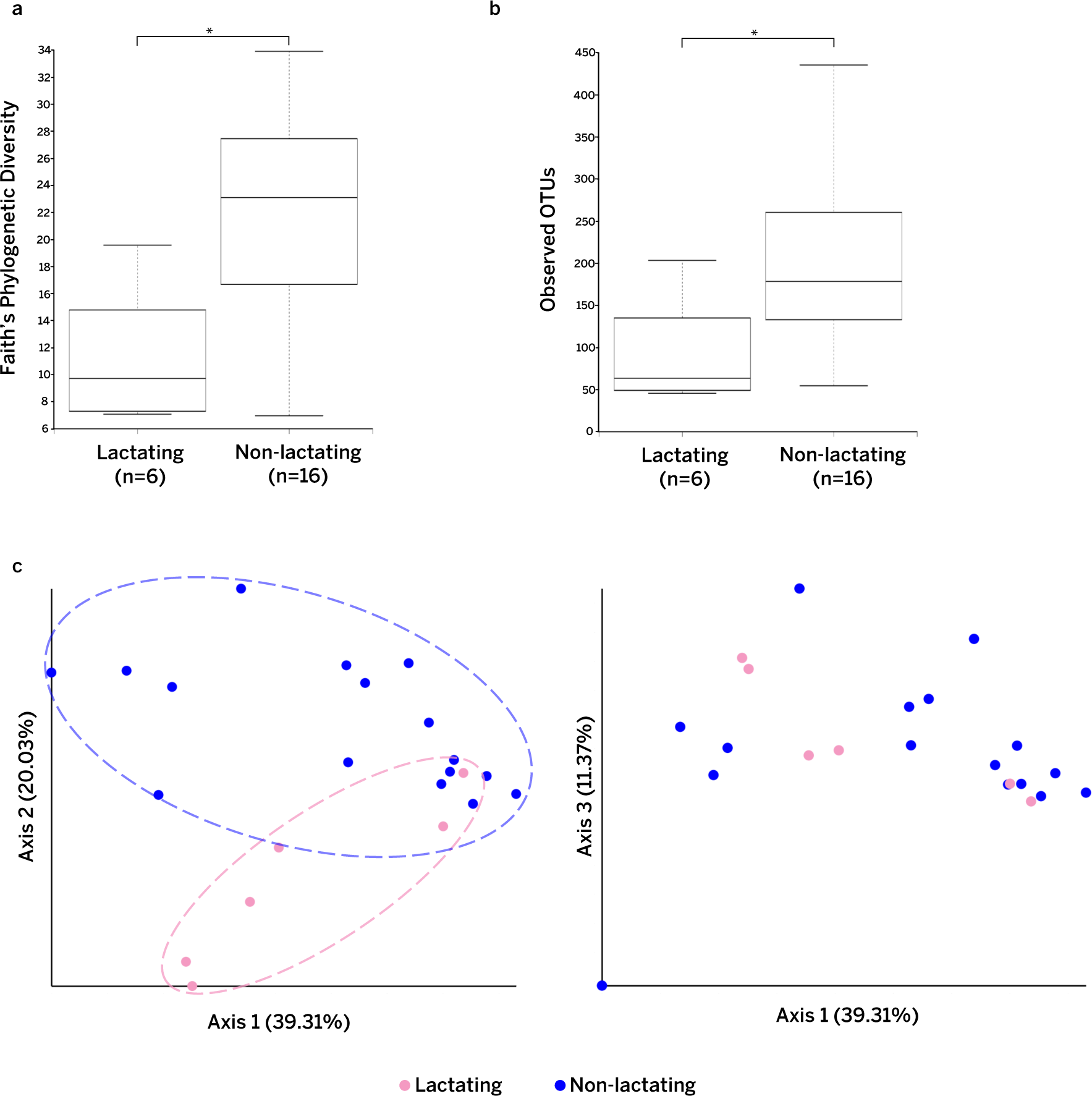
Comparison of microbial diversity in the lactating and non-lactating short-beaked echidna pseudo-pouch. **1a)** Phylogenetic diversity (Faith’s PD) and 1b) species richness (observed OTUs) within lactating and non-lactating pouch samples. **1b)** Principal coordinate analysis (weighted UniFrac) showing clusters of lactating vs non-lactating samples. The left pane shows axes 1 and 2. The right pane shows axes 1 and 3.

Next, we considered whether hormonal changes during breeding season affect the microbiome in the pseudo-pouch. However, the microbiomes of samples within and outside breeding season showed no statistically significant differences for any measure of alpha or beta diversity (Supp Fig S2; p > 0.05). This result also confirmed that the changes seen in lactating animals were independent of breeding season effects.

To better understand the compositional changes observed during lactation, we investigated the taxonomic composition of the different groups. Although all pseudo-pouch groups were dominated by taxa from the same three phyla – Actinobacteria, Proteobacteria, and Firmicutes - non-lactating samples primarily contained Actinobacteria (58%), whereas in the lactating pseudo-pouch, there was a fairly even split between Actinobacteria (42%) and Firmicutes (47%) (Figure 2a). The increase of Firmicutes taxa in the lactating group was found to be statistically significant (ANCOMBC; q = 0.049). Additionally, the lactating pseudo-pouch exhibited a lower taxonomic diversity with only three phyla constituting 98% of observed bacteria.

**Figure 2:**
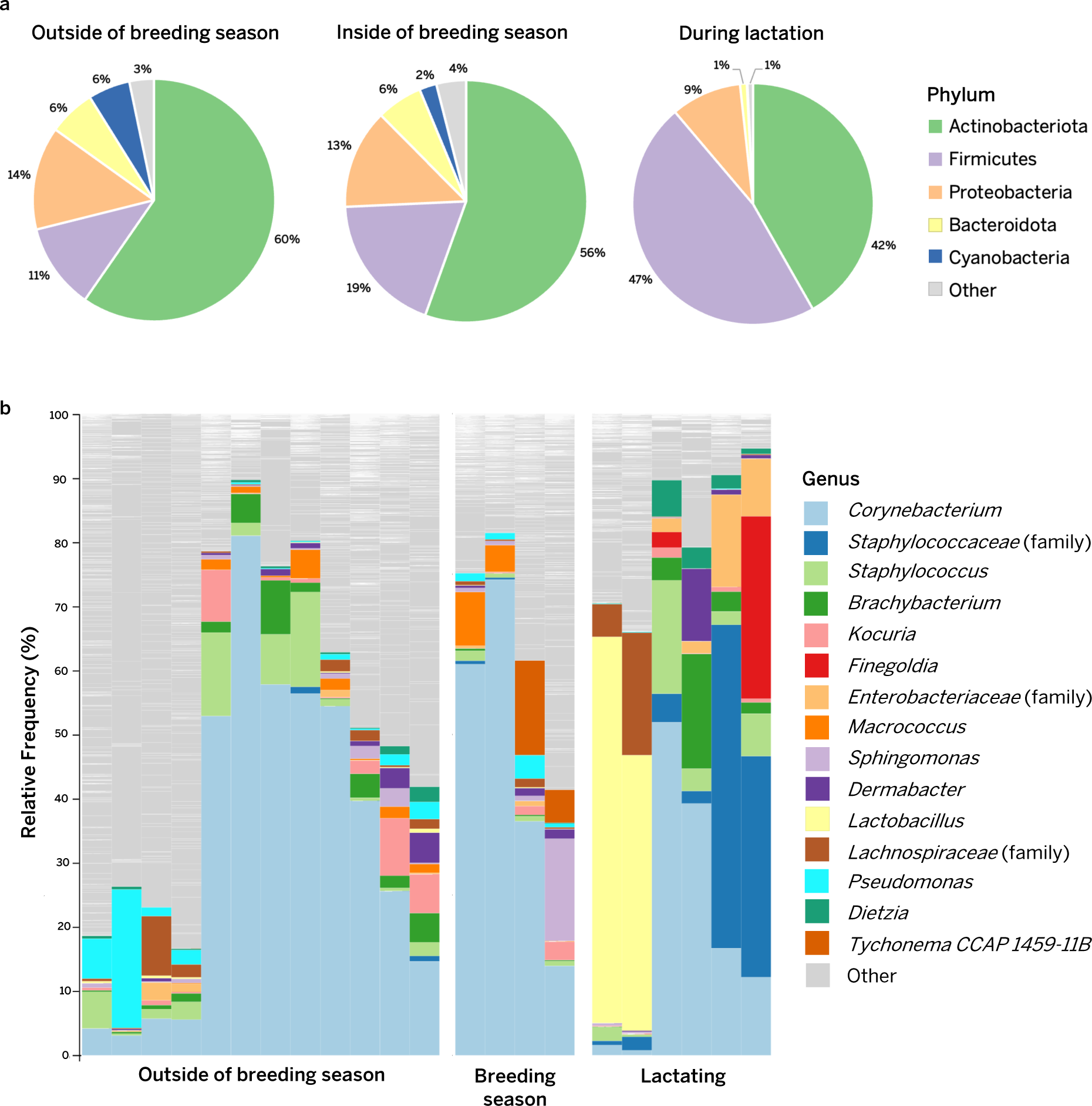
Taxonomy of the short-beaked echidna pseudo-pouch microbiome, comparing bacterial composition inside/outside of breeding season and during lactation. 2a) Relative proportions of bacteria displayed at the phylum level, averaged across sample groups. 2b) Relative proportions of bacteria displayed at the genus level. Top 15 most prevalent taxa are listed to the right of the plot.

In order to gain a more detailed understanding of the differences between the lactating and non-lactating pseudo-pouch microbiome, the samples were further analysed at the genus level (Figure 2b). We observed substantial variation in diversity between individuals. Of the 22 pseudo-pouch samples, 10 were occupied predominantly (i.e. > 50%) by a single bacterial genus (typically *Corynebacterium* spp.). A further 6 samples had relatively even levels of a small number of bacterial taxa. The remaining 6 samples were highly diverse with no apparent dominant taxa. Non-lactating pseudo-pouch samples were predominantly populated by *Corynebacterium* (48.3% in breeding, 37.0% in non-breeding), though some samples were highly diverse with no clear dominant species. The second most prevalent taxon within breeding season was *Sphingomonas* (4.2%) whereas outside of breeding season it was *Staphylococcus* (4.6%). The taxonomic composition of the lactating pseudo-pouch microbiome was substantially altered. The proportion of *Corynebacterium* was greatly reduced (42.6% in non-lactating vs 26.3% in lactating), whereas a number of other taxa increased in abundance, including *Staphylococcaceae* (0.21% in non-lactating vs 20.64% in lactating), *Enterobacteriaceae* (0.69% in non-lactating vs 6.08% in lactating), and *Lactobacillus* (0.13% in non-lactating vs 5.22% in lactating). Interestingly, an unspecified *Finegoldia* species was the third most abundant taxon in lactating individuals, while in the non-lactating pseudo-pouch, it was present only in trace amounts. In general, samples from lactating echidnas exhibited relative increases in proportions of Gram-positive bacteria such as *Staphylococcus*.

### The pseudo-pouch microbiome is a distinct microbial niche

Next, we tested the extent to which other bodily niches and/or the animal’s habitat contributed to changes in the pseudo-pouch microbiome. This included the cloaca, due to the egg passing through this area before being laid into the pseudo-pouch; the mouth, due to the licking behaviour of the dam towards the puggle; dirt from echidna sampling locations (environment), due to the potential exposure of the pseudo-pouch and the milk patch to the ground; and extraction and PCR negative controls, to ensure that any microbes introduced from the laboratory environment were not assumed to be sourced from the animal or its habitat.

**Figure 3:**
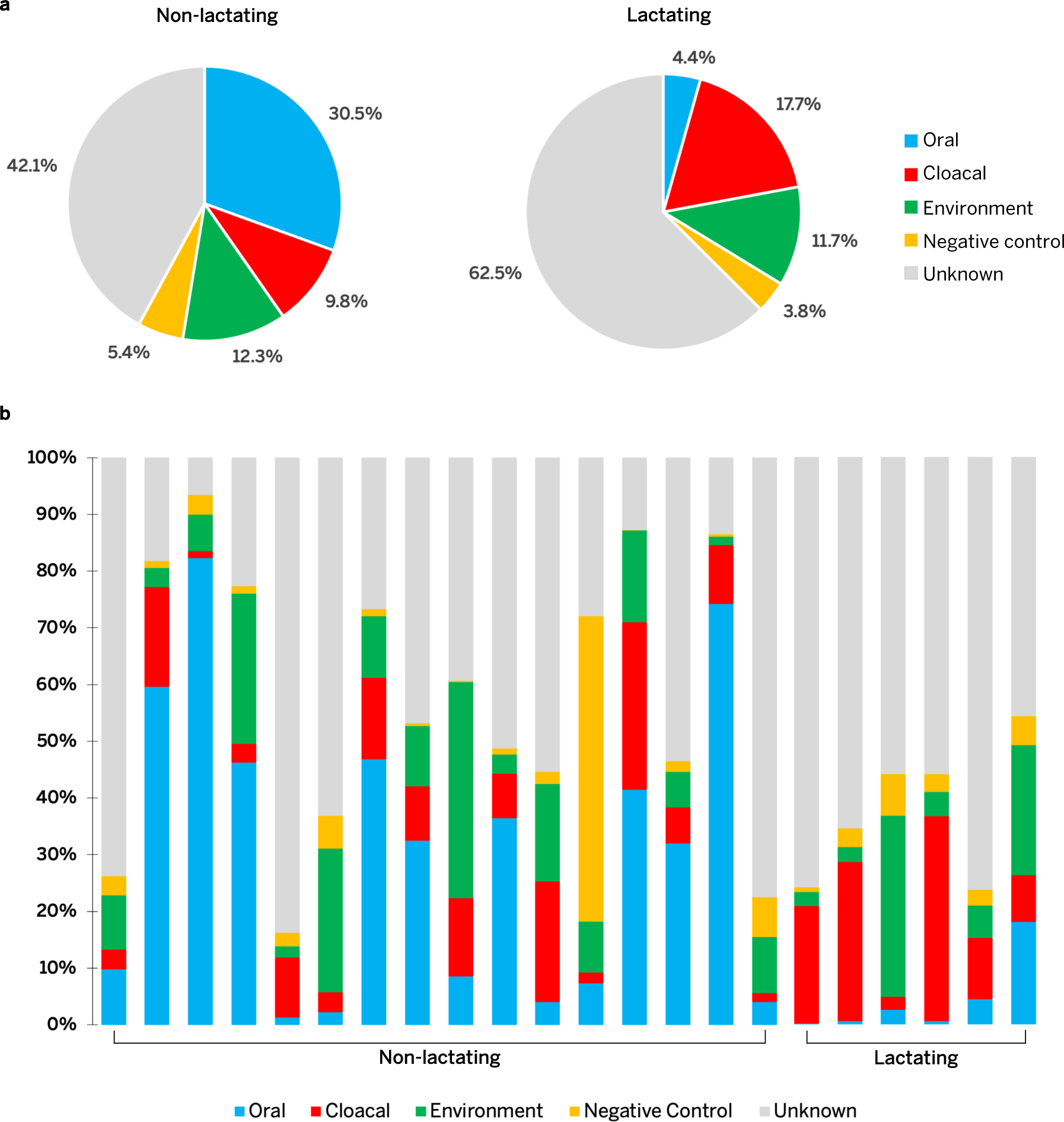
Investigating the origin of the pseudo-pouch microbiome (Sourcetracker2). **3a)** Pie chart showing the average contribution (%) of bacteria from the cloaca, mouth, environment, and negative controls to the pseudo-pouch microbiome in non-lactating and lactating animals. **3b)** Bar graph showing the relative contribution of sources in each pseudo-pouch sample, sorted according to lactation status.

In order to see if any one source group was more similar to the pseudo-pouch than another, we performed beta diversity analysis (weighted UniFrac). This showed that all source groups were significantly different from the pseudo-pouch regardless of lactation status (Supp Fig S3; p < 0.05). We then used Sourcetracker2, a program designed to evaluate the occurrence of microbial seeding between different niches. Taxa derived from the mouth decreased substantially from 30.5% in non-lactating samples to 4.4% in lactating samples. Conversely, taxa derived from the cloaca showed a modest increase (7.9%) during lactation. For both groups, pseudo-pouch taxa were found to be derived primarily from unknown sources, and this proportion increased by 20.4% during lactation. This suggests that the echidna pseudo-pouch microbiome, particularly during lactation, is a specialised microbial niche. The proportion of taxa derived from negative controls (i.e. lab contamination) was relatively small for each group.

We then examined the taxonomic composition of the bacteria contributed by each source group (Supp Fig S4 & S5). Taxa sourced from the cloaca were evenly split between Actinobacteria (primarily *Corynebacterium* spp.) and Firmicutes in non-lactating samples. In lactating samples, the cloacal contribution contained far more Firmicutes, particularly *Lactobacillus* spp. The composition of environment-derived bacteria was relatively unchanged between lactating and non-lactating echidnas. Oral-derived taxa were predominantly *Corynebacterium* spp. for both lactating and non-lactating echidna samples, and lactating samples showed an increase in Proteobacteria. Bacteria derived from unknown sources showed a higher proportion of Firmicutes (particularly *Lactobacillus* spp.) in lactating echidnas.

### Captivity has no significant impact on pseudo-pouch microbiome composition or diversity

Despite improvements in captive management and breeding, the survival rate of pouch young can be low. To explore whether this may be associated with an altered microbiome in captive animals, we compared the microbial diversity in the pseudo-pouch between wild and captive echidnas. Samples collected during lactation were excluded due to a lack of samples from wild individuals. Surprisingly, no significant differences were found between samples derived from wild and captive pseudo-pouches for any of our tested metrics of alpha diversity (Faith’s phylogenetic diversity, richness, evenness) (Supp Fig S6; p > 0.05). Similarly, beta diversity was not significantly different between the groups (Fig 4; weighted UniFrac distance p = 0.571). Overall, our evidence suggests that captivity does not have a significant impact on microbial diversity or composition of the echidna pseudo-pouch microbiome.

**Figure 4:**
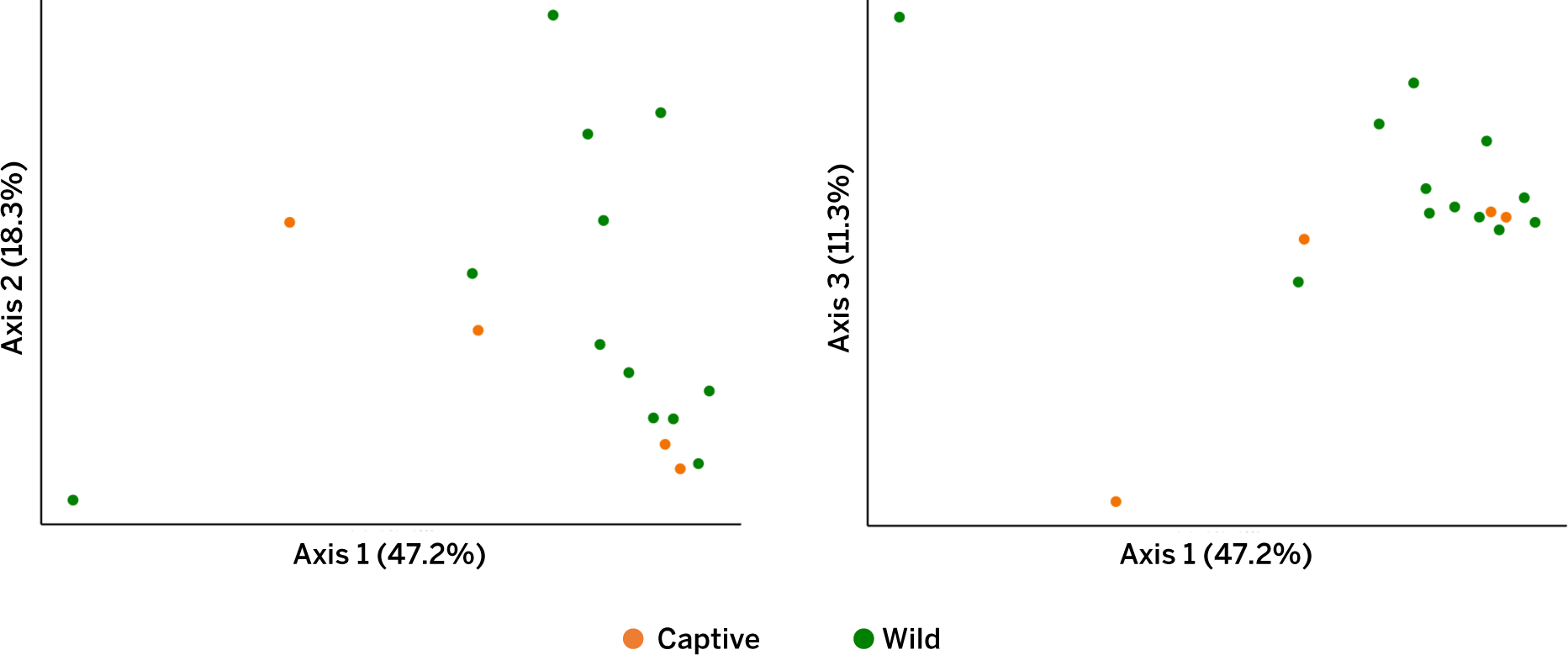
PCoA plot (weighted UniFrac) showing the pseudo-pouch microbiome composition of wild and captive short-beaked echidnas. The left pane shows axes 1 and 2. The right pane shows axes 1 and 3.

## Discussion

The unique biology of monotremes provides invaluable information about the role of lactation in shaping the microbial environment of young lacking functional adaptive immunity^[21]^. In this study we investigated for the first time how the echidna pseudo-pouch microbiome changes during the breeding cycle and lactation, and whether captivity influences the pseudo-pouch microbiome.

In comparison to non-lactating animals, the pseudo-pouches of lactating echidnas exhibited a reduction in microbial richness and phylogenetic diversity. This suggests that only certain bacterial taxa were permitted to remain in the pseudo-pouch during the critical lactation period. For example, during lactation, the proportion of Actinobacteria was significantly reduced whereas Firmicutes were able to proliferate. This contrasts with previous studies of marsupial pouches during lactation, including the quokka, tammar wallaby and southern hairy-nosed wombat, where Actinobacteria made up the vast majority (>80%) of all microbial diversity^[13,22,23]^. This difference may be due to monotreme oviparity and the passage of the egg through the cloaca. Our analysis suggests that the interaction between the cloaca and pseudo-pouch microbiomes is altered during lactation, both in terms of the proportion and taxonomy of bacteria. The proportion of pseudo-pouch bacteria sourced from the cloaca was greater during the lactation period than in non-lactating animals, potentially due to the egg transferring cloacal microbes into the pseudo-pouch. Furthermore, during lactation, most taxa sourced from the cloaca were Firmicutes spp., whereas in non-lactation the majority were Actinobacteria spp. It is possible that the broad increase in Firmicutes and reduction in Actinobacteria seen in the lactating pseudo-pouch are the result of exchange with the cloacal microbiome after the egg is laid.

At the genus level, the echidna pseudo-pouch exhibited decreased *Corynebacterium* and increased *Staphylococcaceae* species during lactation. These genera contain species commensal to human skin, and *Corynebacteria* have been identified as a major constituent of the lactating southern hairy-nosed wombat pouch microbiome^[13]^. However, both genera also contain pathogens such as *C. diphtheriae* and *S. aureus*. In our study, these taxa could only be identified to the genus level, and it therefore remains unknown whether the *Corynebacterium* or *Staphylococcaceae* species that showed a reduced abundance were helpful or harmful. Surprisingly, proportions of *Enterobacteriaceae* spp. were greater in the lactating as compared to the non-lactating pseudo-pouch. This family contains many pathogenic species including *Escherichia coli*, *Salmonella*, and *Shigella*. However, increases in *Enterobacteriaceae* have also been observed in the Tasmanian devil pouch^[13]^, suggesting that these bacteria may be commensals in the pouches of native mammals.

As expected, many of the taxa that became more abundant in the lactating pseudo-pouch were lactic acid bacteria. These bacteria are considered probiotic as they often provide protection by outcompeting pathogens^[23–25]^ and provide a healthy foundation for the infant gastrointestinal microbiome. The proliferation of such commensal species may represent a transition into a more hospitable setting for pouch young development^[13]^.

The shift that we observed between lactating and non-lactating echidna pseudo-pouches is consistent with research in marsupial species. Tammar wallabies undergo broad changes in their pouch microbiomes leading up to and after birth, which involve reductions in overall bacterial richness and Gram-negative species in particular^[22,24]^. A similar phenomenon has been observed in the Tasmanian Devil, where the pouch exhibits reduced abundances of potential pathogens during lactation^[25]^. One proposed mechanism for these changes is skin secretions. These secretions contain antimicrobial peptides including lysozyme, dermcidin, and cathelicidins, the levels of which have been shown to change during different reproductive stages^[14]^. Lysozyme has been shown to bring about decreases in Firmicutes and increases in Bacteroides species in the humans gastrointestinal tract^[26]^ – though this is the opposite of what we observed in the lactating echidna pseudo-pouch. Cathelicidins may work to specifically target pathogens while not affecting commensals, thus bringing about a beneficial microbial environment for pouch young development^[25]^. Cathelicidins have been found in monotremes, raising the possibility that these peptides are key to shaping the pseudo-pouch microbiome during lactation^[17]^. Another potential mechanism of pouch microbiome alteration is that of the milk itself. Marsupial and monotreme milk contains a range of antimicrobial agents including immunoglobulin, lysozyme, transferrin and host defence peptides^[27]^. In marsupials, these antimicrobials are established to protect the infant from infection after ingestion, but may also promote a beneficial microbial environment within the pouch^[14]^. Echidna milk also contains the monotreme-specific peptides MLP and EchAMP, which have demonstrated species-specific activity against a number of both Gram-positive and Gram-negative bacteria^[28,29]^. Currently, we do not understand how these antimicrobial compounds impact the monotreme pseudo-pouch microbiome.

Surprisingly, neither the diversity nor the composition of the pseudo-pouch microbiome changed between captive and wild individuals. There is very little research into the effects of captivity on pouch microbiomes in any species – however, this result contrasts with observations of the Tasmanian devil pouch^[30]^, as well as the gut microbiomes of many other mammals including echidnas, where captivity brought about decreased diversity and major microbial changes^[31,32]^.

The importance of reproductive microbiota in infant immunity is well documented in humans^[1,33]^ but has not been investigated in marsupials or monotremes. Captive-bred puggles face several health challenges not found in wild echidnas, including high coccidia burden leading to secondary yeast infections^[34]^. Our observation that there are minimal differences between the pseudo-pouch microbiomes of wild and captive echidnas suggests that other factors contribute to these health issues.

We considered whether the microbiota of other body parts contribute to the echidna pseudo-pouch microbiome. In particular, we expected that cloacal microbes would be transferred into the pseudo-pouch during oviposition. Oral microbes (derived from pouch-licking behaviour) and microbes from an animal’s environment would also be expected to contribute to the pseudo-pouch microbiome. However, our results show that the pseudo-pouch is highly distinct, particularly during lactation where 62.5% of microbes could not be assigned a specific source. This is consistent with research comparing marsupial milk/pouch samples to other body regions^[13,30]^, though the results may not be directly comparable as these studies did not use specific source-tracking methods. Bacteria derived from the echidnas’ environment stayed relatively consistent between lactating and non-lactating animals, both in terms of relative proportion and taxonomic composition. This suggests that soil microbes are not of concern for the health of pouch young. Our hypothesis that the licking behaviour of echidna dams would increase the proportion of oral-derived microbes in the pseudo-pouch during lactation was not supported; in fact, the oral contribution was substantially decreased during lactation. This contrasts with theories of marsupial development which posit that licking plays a significant role in creating an altered pouch environment for the neonate^[13,35]^. As previously discussed, the cloacal contribution to the pseudo-pouch microbiome was increased during lactation. This result is consistent with what has been observed in Tasmanian devils, where lactating pouches contained more gut-derived bacteria than non-lactating pouches, presumably due to fecal contamination from the pouch young^[25]^. A large proportion of the cloaca-derived bacteria were Lactobacillus spp., which are recognized in other mammals as key vaginal microbiota which colonise the neonate gut and are important for early digestive function^[1,36]^. Together, this suggests that contamination from the digestive tract in the pseudo-pouch is not a cause for concern and may in fact provide beneficial taxa to the flora of the lactating pseudo-pouch and the developing neonate.

### Conclusion

Changes in pouch microbiota associated with lactation have been reported in various mammals and are important for the development and protection of neonates. Monotremes are the only egg-laying mammals but share with marsupials an extended lactation period that commences at a much earlier stage in development. This first characterisation of the short-beaked echidna pseudo-pouch microbiome has revealed changes in diversity and composition during lactation, similar to what has been observed in marsupials. Surprisingly, neither breeding season nor captivity resulted in changes to the pseudo-pouch microbiome. Overall, our results suggest that the pseudo-pouch microbiome is highly distinct and is shaped by lactation rather than by microbes from the environment or other regions of the echidna body. Importantly, we observed that captivity does not have a significant impact on the pseudo-pouch microbiome and is therefore unlikely to contribute to captivity-related health issues in infant echidnas. A better understanding of the molecular mechanisms by which lactation alters the pseudo-pouch microbiome will provide important insights into early echidna development and provide potential candidates for biomedical translation.

## Materials and methods

### Sample collection

This study was carried out in accordance with requirements of the Taronga Animal Ethics Committee (AEC) for the “Collection of opportunistic samples for researchers from live animals during veterinary procedures or routine husbandry procedures” (AEC 4a/02/18), and “Glucose levels and fibre fermentation in monotremes” (AEC 3a/08/21). An overview of the major types of samples used in this study is provided in Table 1; see Supplementary files for full sample metadata. The majority of swabs were collected from captive echidnas at Taronga Zoo, NSW. Further swabs were taken from wild individuals in the care of Taronga Zoo Wildlife Hospital, NSW, as well as road-kill echidnas from Kangaroo Island (collected by Dr Peggy Rismiller) and the Adelaide Hills (sourced through EchidnaCSI). Pouch swabs were taken from the pseudo-pouch in lactating animals and the corresponding abdominal region in non-lactating animals; oral swabs were taken from inside the beak; cloacal swabs were taken from the external cloacal opening and the internal cloaca; and environment samples were taken from dirt within captive echidnas’ enclosures, or from the ground next to deceased wild echidnas. Where swabs were collected within the lab, a lab negative control swab of the air was also taken for downstream identification of contaminants. All swabs were frozen at -70°C to prevent DNA degradation.

### DNA extraction

All extractions took place in a still air class II biosafety cabinet which was decontaminated with 10% bleach. Total genomic DNA was extracted from swab samples using the ZymoBIOMICS^TM^ DNA Miniprep Kit according to the manufacturer’s instructions, except that buffers were aliquoted prior to the introduction of samples into the hood in order to minimise contamination. In order to account for laboratory-based contamination, one extraction blank control in which no swab was added to the lysis tube was included with each set of extractions (total number of extraction blank controls = 10). Scissors used to cut swab tips into tubes were wiped on bleach-soaked paper towel in between each sample to avoid cross-contamination. Bead-beating step was carried out using a Disruptor Genie® cell disruptor for 15 minutes. DNA samples were stored at -20 °C until required for further use.

### Amplicon library preparation and quantification

The V4 region of the bacterial 16S gene was amplified in all samples using PCR. Additional negative (no DNA added; n = 10) and positive (DNA known to amplify successfully; from short beaked echidna scat sample) controls were included in each set of PCR reactions. Each sample was amplified using the 515F forward primer: 5’-AATGATACGGCGACCACCGAGATCTACACTATGGTAATTGTGTGCCAGCGCCG CGGTAA-3’, and the uniquely barcoded 806R reverse primer: 5’-CAAGCAGAAGACGGCATACGAGAT-nnnnnnnnnnnn-AGTCAGTCAGCCGGACTACGGGTTCTAAT-3’, with the 12 n’s representing a unique barcode sequence for downstream identification.

Reactions containing 18.7 μL H_2_O, 2.5 μL High Fidelity buffer, 1 μL 50 mM MgSO_4_, 0.2 μL 100 mM dNTP mix, 0.5 μL 10 μM forward primer, 0.1 μL Platinum™ Taq DNA Polymerase High Fidelity (Thermofisher), 1 μL sample DNA and 1 μL 5 μM unique reverse primer were amplified using an initial denaturation step at 94 °C for 3 minutes, followed by 35 cycles of denaturation at 94 °C for 45 seconds, annealing at 50 °C for 60 seconds, and elongation at 68 °C for 90 seconds, with a final adenylation step at 68 °C for 7 minutes. In order to assess whether the desired fragment of DNA (∼390 bp) had been successfully amplified, gel electrophoresis was performed using a 1.5% agarose gel with RedSafe™ Nucleic Acid Staining Solution (20,000x) (iNtRON Biotechnology) run at 100 V for 30 minutes.

2 μL of each sample (excluding PCR positive controls) was sent to the Australian Cancer Research Facility (ACRF) for fluorometric quantification using a Qubit™. Samples were then pooled to equimolar concentration and cleaned using the Agencourt AMPure XP PCR purification system (Beckman Coulter) according to the manufacturer’s instructions, before being sent to ACRF for a final quantification and quality assessment using an Agilent 2100 Bioanalyzer system. The samples were pooled to 4 nM and sequenced on an Illumina MiSeq (v2, 2 x 250 bp 15M reads) at ACRF.

### Data analysis

Demultiplexed reads were quality-controlled and adaptor-trimmed using the FASTQ pre-processor program fastp^[37]^. The sequences were further processed and analysed using the QIIME2 (v. 2020.2.0) bioinformatic pipeline^[38]^. Paired-end reads were joined using vsearch^[39]^. Deblur^[40]^ was used to denoise the reads into amplicon sequence variants (ASVs); a trim length of 252bp was used based on visualisation of the joined reads. The prevalence-based method in decontam^[41]^ was used to identify putative contaminants (threshold score 0.5), which were removed from the feature table along with singletons (Supp Fig S7; Supp Fig S8). The taxonomic composition of negative control samples prior to contaminant removal is displayed in Supp Fig S9. The table was rarefied using a maximum depth of 6300.

Diversity analyses were performed using core metrics phylogenetic at a sampling depth of 1000. ASVs were taxonomically classified using the SILVA 16S V4 classifier^[42]^ (version 138 515f 806r). Alpha diversity was estimated using Observed Operational Taxonomic Units (OTUs), Pielou’s evenness, Shannon’s diversity, and Faith’s phylogenetic diversity^[43]^; Beta diversity was estimated using weighted UniFrac distances^[44]^. UniFrac distances were visualised using Principal Coordinate Analysis (PCoA) plots. Significance of alpha and beta diversity was assessed using Kruskal-Wallis pairwise tests and Permutational Multivariate Analysis of Variance (PERMANOVA) tests respectively. Statistical significance of differentially abundant taxa between groups was undertaken using the ANCOM plugin for Qiime2^[45]^. Sourcetracker2^[46]^ (ver. 2.0.1) was used to estimate the contribution of oral, cloacal and environmental bacteria to the bacterial composition of the pouch microbiome. A rarefaction depth of 1000 was applied to all sinks. The per_sink_feature_assignments flag was used to assign QIIME2 feature IDs to each identified source OTU.

## Acknowledgements

I.W. was supported by a University of Adelaide Research Scholarship. The authors acknowledge the staff of Taronga Wildlife Hospital for their role in sample collection.

## Notes

https://github.com/isa-wilson/echidna_pouch_microbiome

### Competing Interest Statement

The authors have declared no competing interest.

## Works Cited

1 Mueller, N. T., Bakacs, E., Combellick, J., Grigoryan, Z. & Dominguez-Bello, M. G. The infant microbiome development: mom matters. Trends Mol. Med. 21, 109–117, 10.1016/j.molmed.2014.12.002 (2015).

2 Palmer, C., Bik, E. M., DiGiulio, D. B., Relman, D. A. & Brown, P. O. Development of the Human Infant Intestinal Microbiota. PLoS Biol. 5, e177, doi:10.1371/journal.pbio.0050177 (2007).

3 Dominguez-Bello, M. G. et al. Delivery mode shapes the acquisition and structure of the initial microbiota across multiple body habitats in newborns. Proceedings of the National Academy of Sciences 107, 11971–11975, doi:10.1073/pnas.1002601107 (2010).

4 Barile, D. & Rastall, R. A. Human milk and related oligosaccharides as prebiotics. Curr. Opin. Biotechnol. 24, 214–219, 10.1016/j.copbio.2013.01.008 (2013).

5 Gopalakrishna, K. P. & Hand, T. W. Influence of Maternal Milk on the Neonatal Intestinal Microbiome. Nutrients 12, 823 (2020).

6 Heijtz, R. D. et al. Normal gut microbiota modulates brain development and behavior. Proceedings of the National Academy of Sciences 108, 3047–3052, doi:10.1073/pnas.1010529108 (2011).

7 Smith, C. L. et al. Identification of a human neonatal immune-metabolic network associated with bacterial infection. Nature Communications 5, 4649, doi:10.1038/ncomms5649 (2014).

8 Gürdeniz, G. et al. Neonatal metabolome of caesarean section and risk of childhood asthma. Eur. Respir. J. 59, 2102406, doi:10.1183/13993003.02406-2021 (2022).

9 Hoang, D. M., Levy, E. I. & Vandenplas, Y. The impact of Caesarean section on the infant gut microbiome. Acta Paediatr. 110, 60–67, 10.1111/apa.15501 (2021).

10 Kuhle, S., Tong, O. S. & Woolcott, C. G. Association between caesarean section and childhood obesity: a systematic review and meta-analysis. Obes. Rev. 16, 295–303, 10.1111/obr.12267 (2015).

11 Sevelsted, A., Stokholm, J., Bønnelykke, K. & Bisgaard, H. Cesarean Section and Chronic Immune Disorders. Pediatrics 135, e92–e98, doi:10.1542/peds.2014-0596 (2015).

12 Tanoey, J., Gulati, A., Patterson, C. & Becher, H. Risk of Type 1 Diabetes in the Offspring Born through Elective or Non-elective Caesarean Section in Comparison to Vaginal Delivery: a Meta-Analysis of Observational Studies. Curr. Diab. Rep. 19, 124, doi:10.1007/s11892-019-1253-z (2019).

13 Weiss, S., Taggart, D., Smith, I., Helgen, K. M. & Eisenhofer, R. Host reproductive cycle influences the pouch microbiota of wild southern hairy-nosed wombats (Lasiorhinus latifrons). Animal Microbiome 3, 13, doi:10.1186/s42523-021-00074-8 (2021).

14 Cheng, Y. & Belov, K. Antimicrobial Protection of Marsupial Pouch Young. Front. Microbiol. 8, doi:10.3389/fmicb.2017.00354 (2017).

15 Peel, E. et al. Marsupial and monotreme cathelicidins display antimicrobial activity, including against methicillin-resistant Staphylococcus aureus. Microbiology 163, 1457–1465, 10.1099/mic.0.000536 (2017).

16 Maidment, T. & Eisenhofer, R. Pouch bacteria: an understudied and potentially important facet of marsupial reproduction. Microbiol. Aust. 44, 41–44, 10.1071/MA23010 (2023).

17 Wang, J. et al. Ancient Antimicrobial Peptides Kill Antibiotic-Resistant Pathogens: Australian Mammals Provide New Options. PLoS One 6, e24030, doi:10.1371/journal.pone.0024030 (2011).

18 Griffiths, M. The Biology of the Monotremes. (Academic Press, 1978).

19 Rismiller, P. D. & Seymour, R. S. The Echidna. Sci. Am. 264, 96–103 (1991).

20 Rismiller, P. D. & McKelvey, M. W. Activity and behaviour of lactating echidnas (Tachyglossus aculeatus multiaculeatus) from hatching of egg to weaning of young. Aust. J. Zool. 57, 265–273, 10.1071/ZO09031 (2009).

21 Comizzoli, P., Power, M. L., Bornbusch, S. L. & Muletz-Wolz, C. R. Interactions between reproductive biology and microbiomes in wild animal species. Animal Microbiome 3, 87, doi:10.1186/s42523-021-00156-7 (2021).

22 Chhour, K. L., Hinds, L. A., Jacques, N. A. & Deane, E. M. An observational study of the microbiome of the maternal pouch and saliva of the tammar wallaby, Macropus eugenii, and of the gastrointestinal tract of the pouch young. Microbiology 156, 798–808, doi:10.1099/mic.0.031997-0 (2010).

23 Charlick, J., Manessis, C., Stanley, N., Waring, H. & Cockson, A. QUANTITATIVE ALTERATIONS OF THE AEROBIC BACTERIAL FLORA OF THE POUCH OF SETONIX BRACHYURUS (QUOKKA) DURING OESTRUS, ANOESTRUS, PREGNANCY AND LACTATING ANOESTRUS (POUCH YOUNG). Aust. J. Exp. Biol. Med. Sci. 59, 743–751, 10.1038/icb.1981.64 (1981).

24 Old, J. M. & Deane, E. M. The effect of oestrus and the presence of pouch young on aerobic bacteria isolated from the pouch of the tammar wallaby, *Macropus eugenii*. Comp. Immunol. Microbiol. Infect. Dis. 21, 237–245, doi:10.1016/s0147-9571(98)00022-8 (1998).

25 Peel, E. et al. Cathelicidins in the Tasmanian devil (Sarcophilus harrisii). Sci. Rep. 6, doi:10.1038/srep35019 (2016).

26 Maga, E. A. et al. Consumption of Lysozyme-Rich Milk Can Alter Microbial Fecal Populations. Appl. Environ. Microbiol. 78, 6153–6160, doi:10.1128/AEM.00956-12 (2012).

27 Stannard, H. J., Miller, R. D. & Old, J. M. Marsupial and monotreme milk—a review of its nutrient and immune properties. PeerJ 8, 1–31, doi:10.7717/peerj.9335 (2020).

28 Bisana, S. et al. Identification and Functional Characterization of a Novel Monotreme-Specific Antibacterial Protein Expressed during Lactation. PLoS One 8, e53686, doi:10.1371/journal.pone.0053686 (2013).

29 Enjapoori, A. K. et al. Monotreme lactation protein is highly expressed in monotreme milk and provides antimicrobial protection. Genome Biol. Evol. 6, 2754–2773, doi:10.1093/gbe/evu209 (2014).

30 Cheng, Y. et al. The Tasmanian devil microbiome—implications for conservation and management. Microbiome 3, doi:10.1186/s40168-015-0143-0 (2015).

31 McKenzie, V. J. et al. The Effects of Captivity on the Mammalian Gut Microbiome. Integr. Comp. Biol. 57, 690–704, doi:10.1093/icb/icx090 (2017).

32 Perry, T. et al. Characterising the Gut Microbiomes in Wild and Captive Short-Beaked Echidnas Reveals Diet-Associated Changes. Front. Microbiol. 13, doi:10.3389/fmicb.2022.687115 (2022).

33 Nyangahu, D. D. & Jaspan, H. B. Influence of maternal microbiota during pregnancy on infant immunity. Clin. Exp. Immunol. 198, 47–56, doi:10.1111/cei.13331 (2019).

34. Tobias, G. in *Current Therapy in Medicine of Australian Mammals* (ed Larry Portas Timothy Vogelnest) Ch. 29, 425–432 (CSIRO Publishing, 2019).

35 Ambatipudi, K., Joss, J., Raftery, M. & Deane, E. A proteomic approach to analysis of antimicrobial activity in marsupial pouch secretions. Dev. Comp. Immunol. 32, 108–120, 10.1016/j.dci.2007.04.009 (2008).

36 Chen, C. et al. The microbiota continuum along the female reproductive tract and its relation to uterine-related diseases. Nature Communications 8, 875, doi:10.1038/s41467-017-00901-0 (2017).

37 Chen, S., Zhou, Y., Chen, Y. & Gu, J. fastp: an ultra-fast all-in-one FASTQ preprocessor. Bioinformatics 34, i884–i890, doi:10.1093/bioinformatics/bty560 (2018).

38 Bolyen, E. et al. Reproducible, interactive, scalable and extensible microbiome data science using QIIME 2. Nat. Biotechnol. 37, 852–857, doi:10.1038/s41587-019-0209-9 (2019).

39 Rognes, T., Flouri, T., Nichols, B., Quince, C. & Mahé, F. VSEARCH: a versatile open source tool for metagenomics. PeerJ 4, e2584, doi:10.7717/peerj.2584 (2016).

40 Amir, A. et al. Deblur Rapidly Resolves Single-Nucleotide Community Sequence Patterns. mSystems 2, e00191–00116, doi:10.1128/mSystems.00191-16 (2017).

41 Davis, N. M., Proctor, D. M., Holmes, S. P., Relman, D. A. & Callahan, B. J. Simple statistical identification and removal of contaminant sequences in marker-gene and metagenomics data. Microbiome 6, 226, doi:10.1186/s40168-018-0605-2 (2018).

42 Quast, C. et al. The SILVA ribosomal RNA gene database project: improved data processing and web-based tools. Nucleic Acids Res. 41, D590–D596, doi:10.1093/nar/gks1219 (2012).

43 Faith, D. P. Conservation evaluation and phylogenetic diversity. Biol. Conserv. 61, 1–10, 10.1016/0006-3207(92)91201-3 (1992).

44 Lozupone, C. & Knight, R. UniFrac: a New Phylogenetic Method for Comparing Microbial Communities. Appl. Environ. Microbiol. 71, 8228–8235, doi:10.1128/AEM.71.12.8228-8235.2005 (2005).

45 Mandal, S. et al. Analysis of composition of microbiomes: a novel method for studying microbial composition. Microb. Ecol. Health Dis. 26, 27663, doi:10.3402/mehd.v26.27663 (2015).

46 Knights, D. et al. Bayesian community-wide culture-independent microbial source tracking. Nature Methods 8, 761–763, doi:10.1038/nmeth.1650 (2011).

