## Supplementary data for "Pouch microbiome changes during lactation in the short-beaked echidna (*Tachyglossus aculeatus*)"

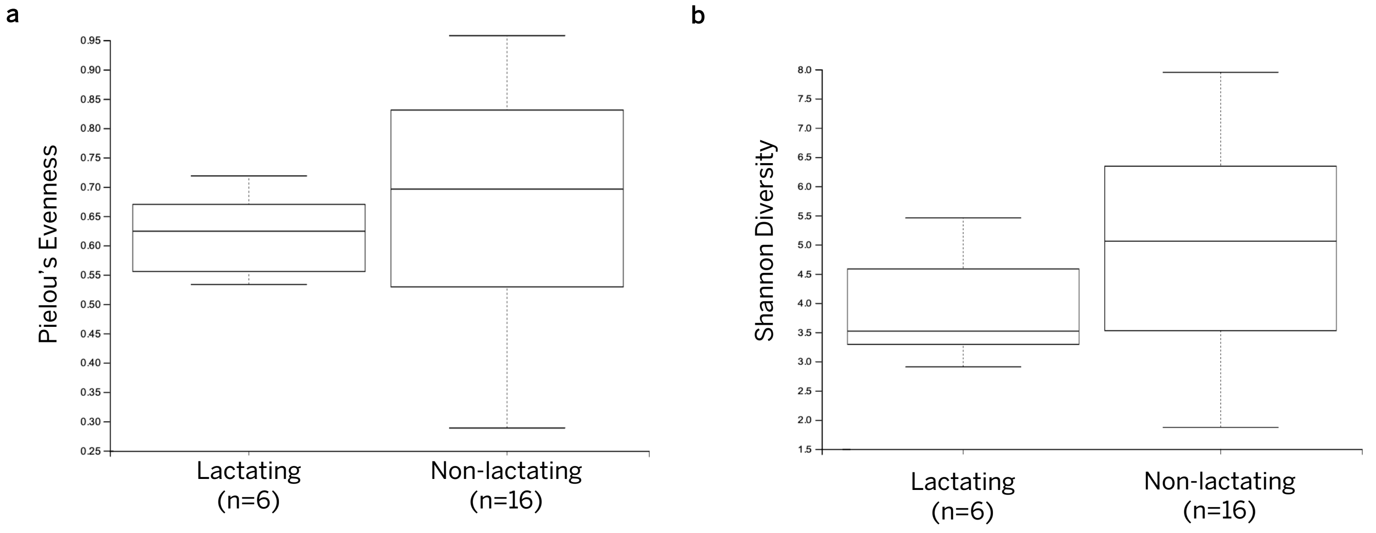


**Supplementary figure S1: Alpha diversity analysis of lactating and non-lactating pseudo-pouch samples**. a) Species evenness measured by Pielou's evenness; b) species diversity measured by Shannon's index.

**Supplementary table S1: statistical significance of alpha diversity analyses on lactating vs non-lactating pseudo-pouch samples.**

| Alpha diversity metric | H | p-value | q-value |
|---|---|---|---|
| Observed OTUs | 5.226 | 0.022 | 0.022 |
| Shannon Diversity | 1.761 | 0.185 | 0.185 |
| Pielou's Evenness | 0.266 | 0.606 | 0.606 |
| Faith's Phylogenetic Diversity | 5.223 | 0.022 | 0.022 |

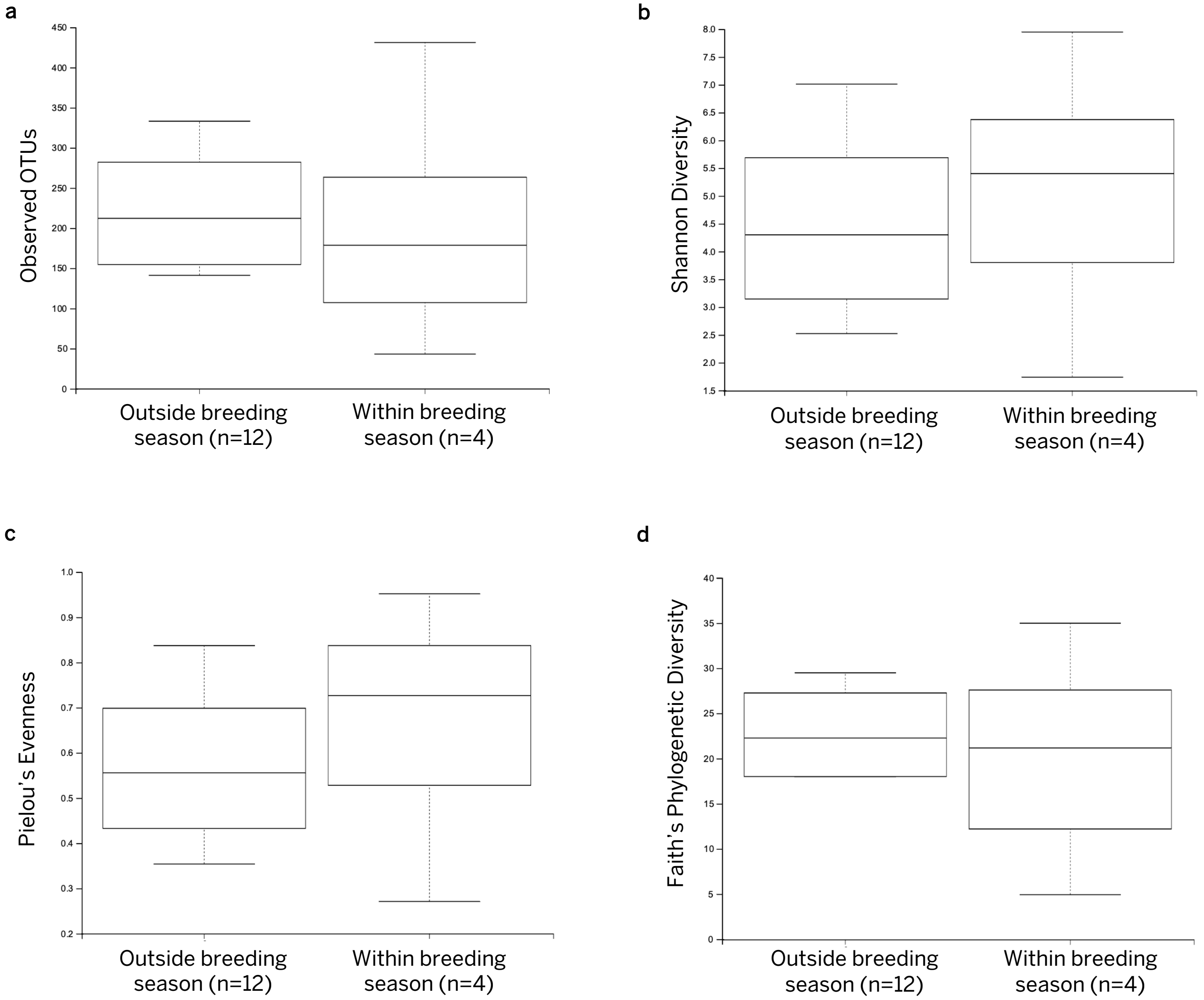


**Supplementary figure S2: Alpha diversity analysis of pseudo-pouch samples taken within vs outside of breeding season. a)** Species richness measured by observed OTUs; **b)** species richness measured by Shannon's index; **c)** species evenness measured by Pielou's evenness; **d)** phylogenetic diversity measured by Faith's PD.

**Supplementary table S2: statistical significance of alpha diversity analyses on samples within breeding season vs outside of breeding season.**

| Alpha diversity metric | H | p-value | q-value |
|---|---|---|---|
| Observed OTUs | 0.132 | 0.716 | 0.716 |
| Shannon Diversity | 0.529 | 0.467 | 0.467 |
| Pielou's Evenness | 0.941 | 0.332 | 0.332 |
| Faith's Phylogenetic Diversity | 0.059 | 0.808 | 0.808 |

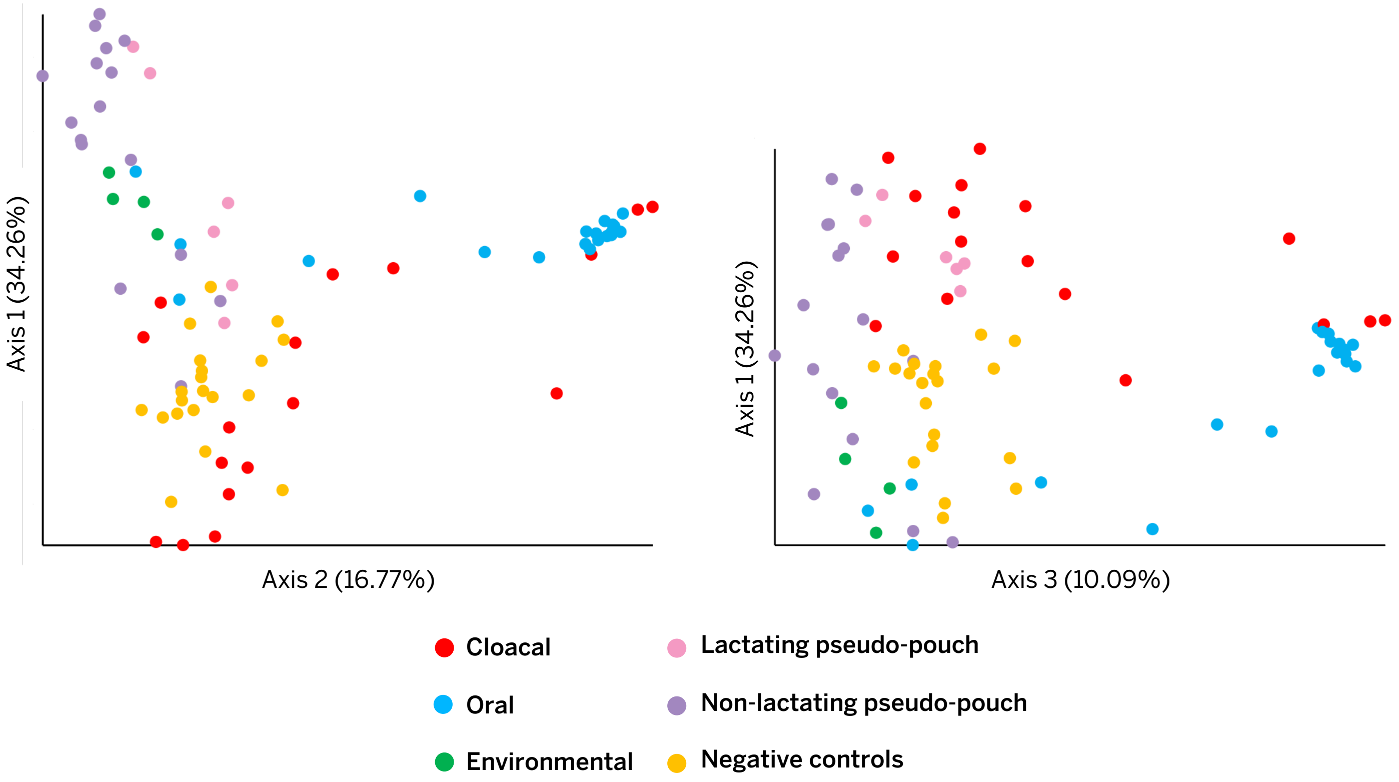


**Supplementary Figure S3: PCoA plot (weighted UniFrac) showing the compositional similarity between the pseudo-pouch microbiome and three potential sources of its microbial diversity (cloaca, mouth, environment) plus negative controls**. The left pane shows axes 1 and 2. The right pane shows axes 1 and 3.

**Supplementary Table S3: Statistical significance of beta diversity analysis (weighted UniFrac) on pouch microbiome samples and their proposed sources (cloaca, mouth, environment, negative controls).**

| Group 1 | Group 2 | Sample Size | Permutations | pseudo-F | p-value | q-value |
|---|---|---|---|---|---|---|
| Non-lactating pseudo-pouch | Cloacal | 33 | 999 | 13.025 | 0.001 | 0.002 |
| | Oral | 36 | 999 | 27.245 | 0.001 | 0.002 |
| | Environment | 20 | 999 | 2.507 | 0.036 | 0.049 |
| | Extraction blank | 26 | 999 | 9.371 | 0.001 | 0.002 |
| | PCR negative | 26 | 999 | 9.781 | 0.001 | 0.002 |
| Lactating pseudo-pouch | Cloacal | 23 | 999 | 3.798 | 0.004 | 0.007 |
| | Oral | 26 | 999 | 11.345 | 0.001 | 0.002 |
| | Environment | 10 | 999 | 3.509 | 0.005 | 0.008 |
| | Extraction blank | 16 | 999 | 6.541 | 0.001 | 0.002 |
| | PCR negative | 16 | 999 | 6.579 | 0.001 | 0.002 |

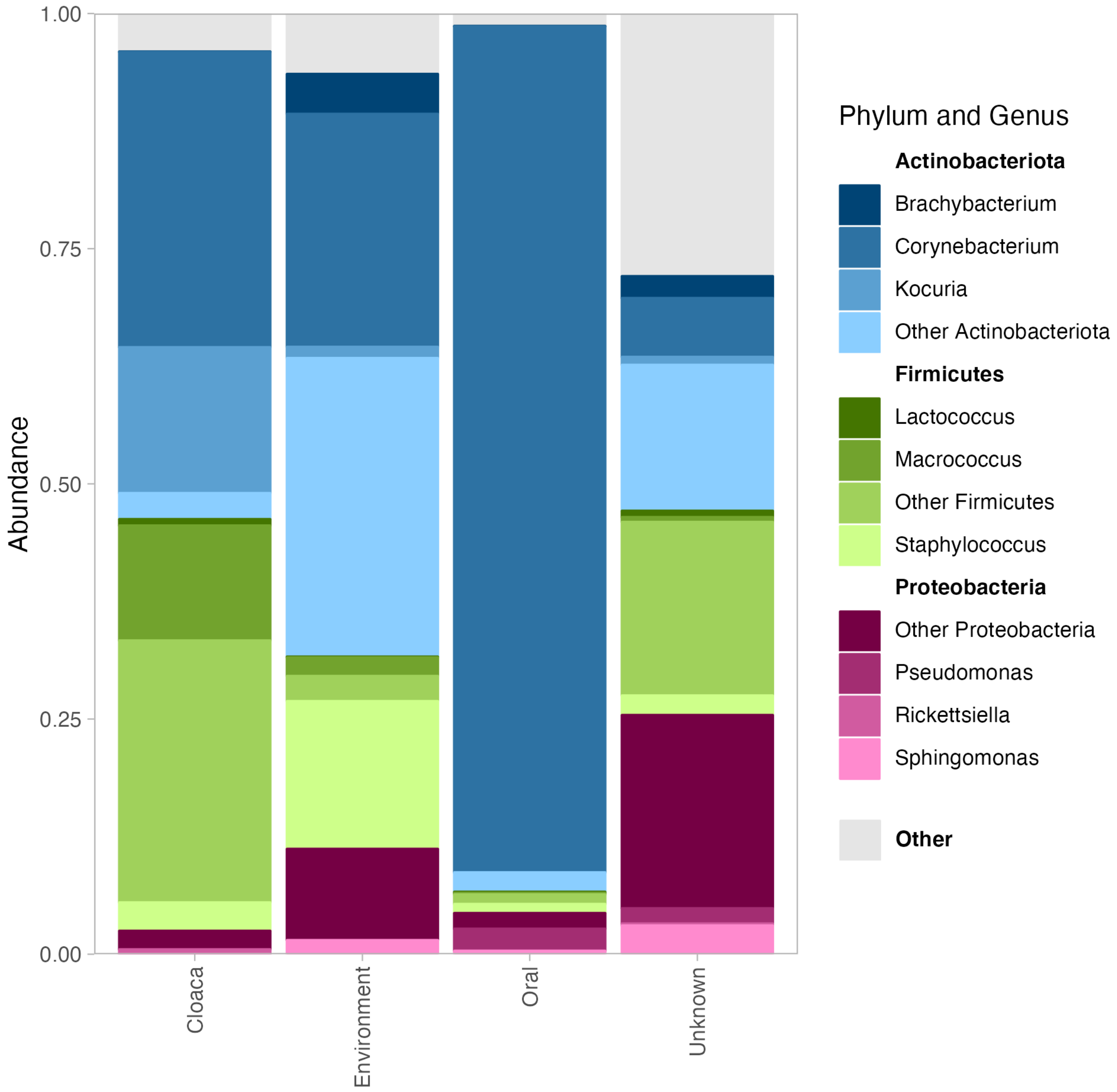


**Supplementary figure S4: Relative abundance chart showing the taxonomic composition of non-lactating echidna pseudo-pouch microbes derived from the cloaca, environment, mouth, and unknown sources.** Taxa are displayed at the phylum and genus level. Source taxa were identified using Sourcetracker2. Plot was generated using the Phyloseq and Fantaxtic R packages.

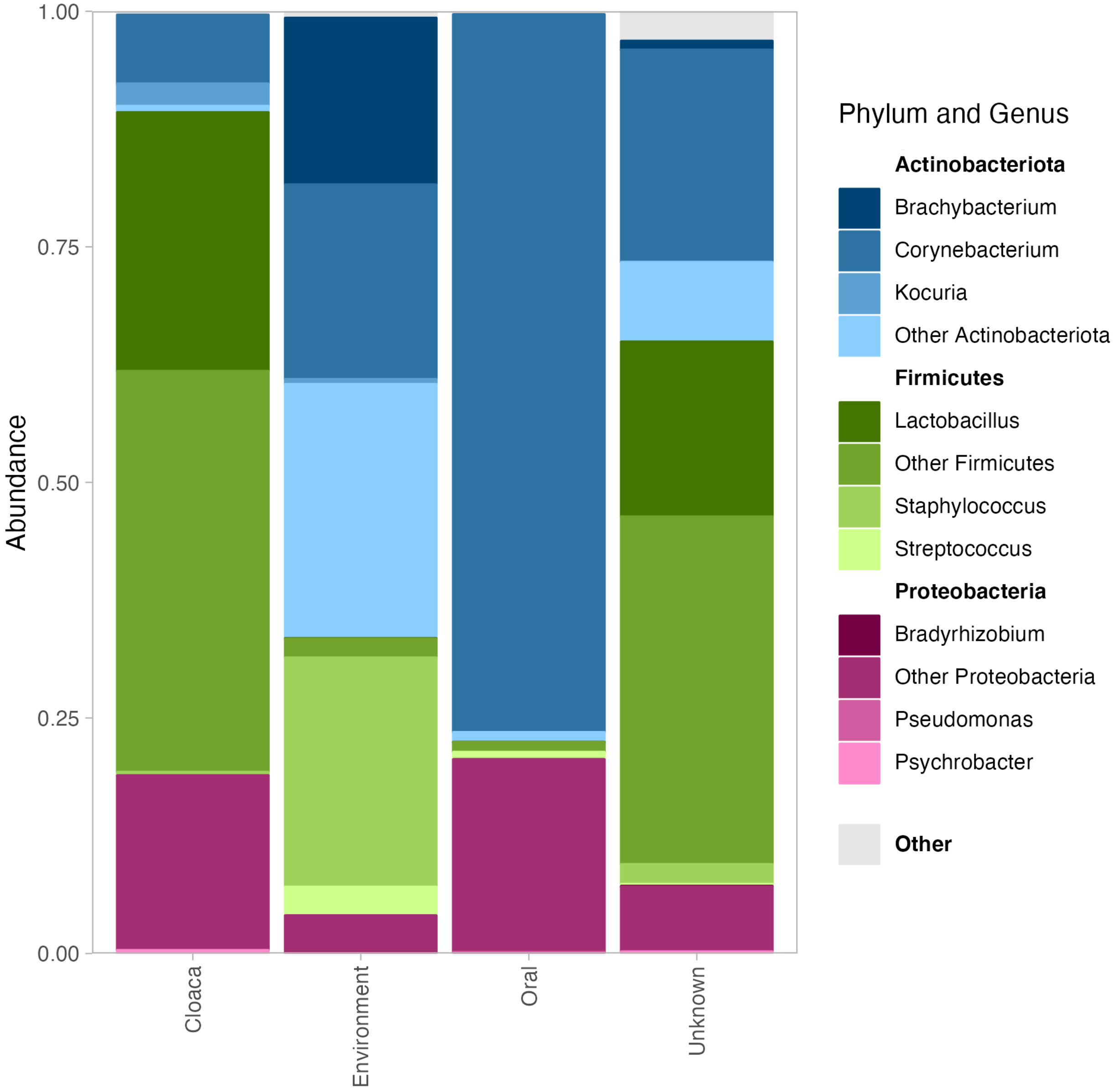


**Supplementary figure S5: Relative abundance chart showing the taxonomic composition of lactating echidna pseudo-pouch microbes derived from the cloaca, environment, mouth, and unknown sources.** Taxa are displayed at the phylum and genus level. Source taxa were identified using Sourcetracker2. Plot was generated using the Phyloseq and Fantaxtic R packages.

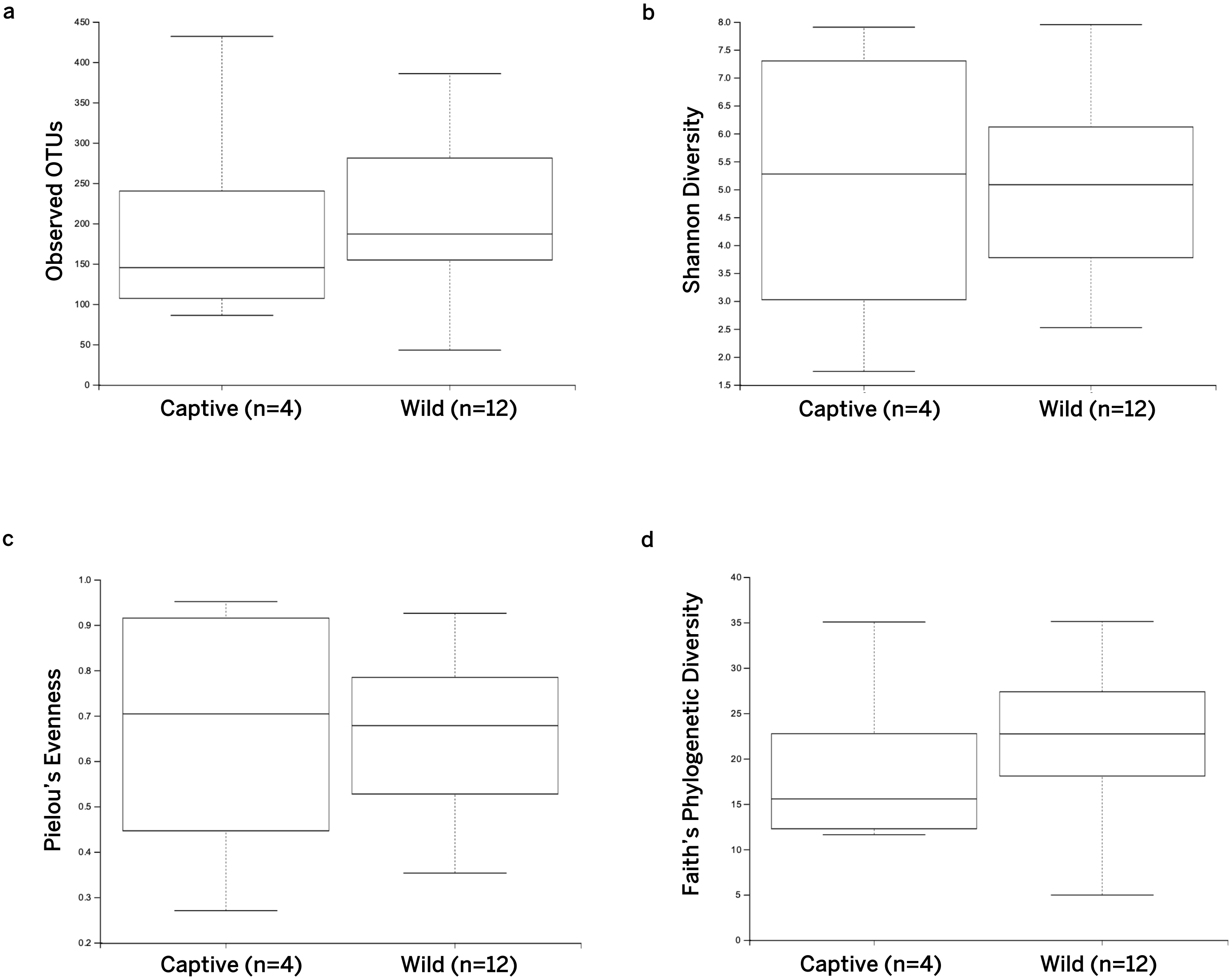


**Supplementary figure S6: Alpha diversity analysis of captive and wild non-lactating pseudo-pouch samples. a)** Species richness measured by observed OTUs; **b)** species richness measured by Shannon's index; **c)** species evenness measured by Pielou's evenness; **d)** phylogenetic diversity measured by Faith's PD.

**Supplementary table S4: Statistical significance of alpha diversity analyses on captive and wild non-lactating pseudo-pouch samples.**

| Alpha diversity metric | H | p-value | q-value |
|---|---|---|---|
| Observed OTUs | 0.132 | 0.716 | 0.716 |
| Shannon Diversity | 0.015 | 0.903 | 0.903 |
| Pielou's Evenness | 0.015 | 0.903 | 0.903 |
| Faith's Phylogenetic Diversity | 0.368 | 0.544 | 0.544 |

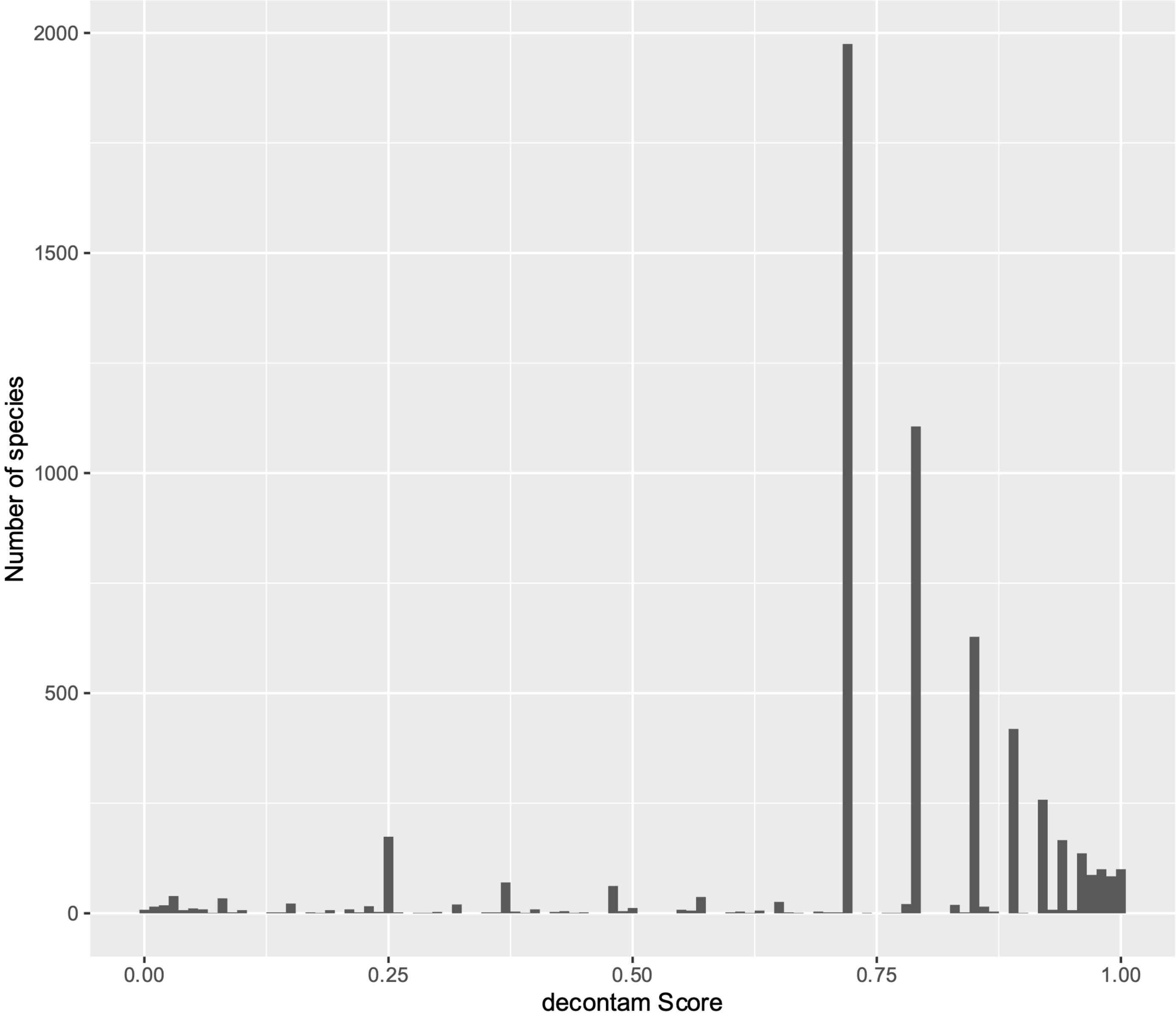


**Supplementary figure S7: Histogram of decontam scores.**

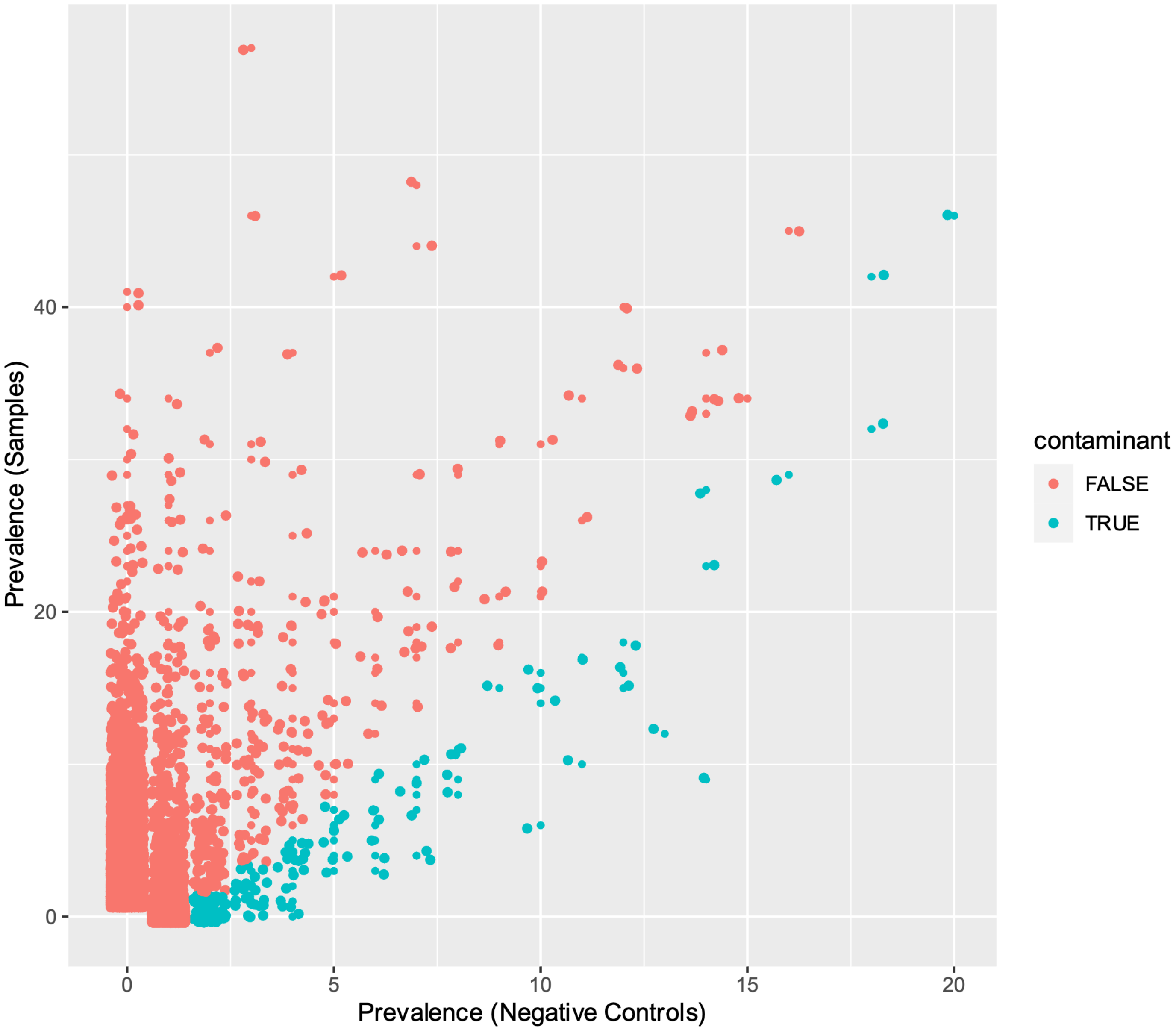


**Supplementary figure S8: Prevalence/prevalence plot of taxa identified in biological samples vs negative controls.** Taxa identified by decontam as contaminants (score >0.5) are shown in blue.

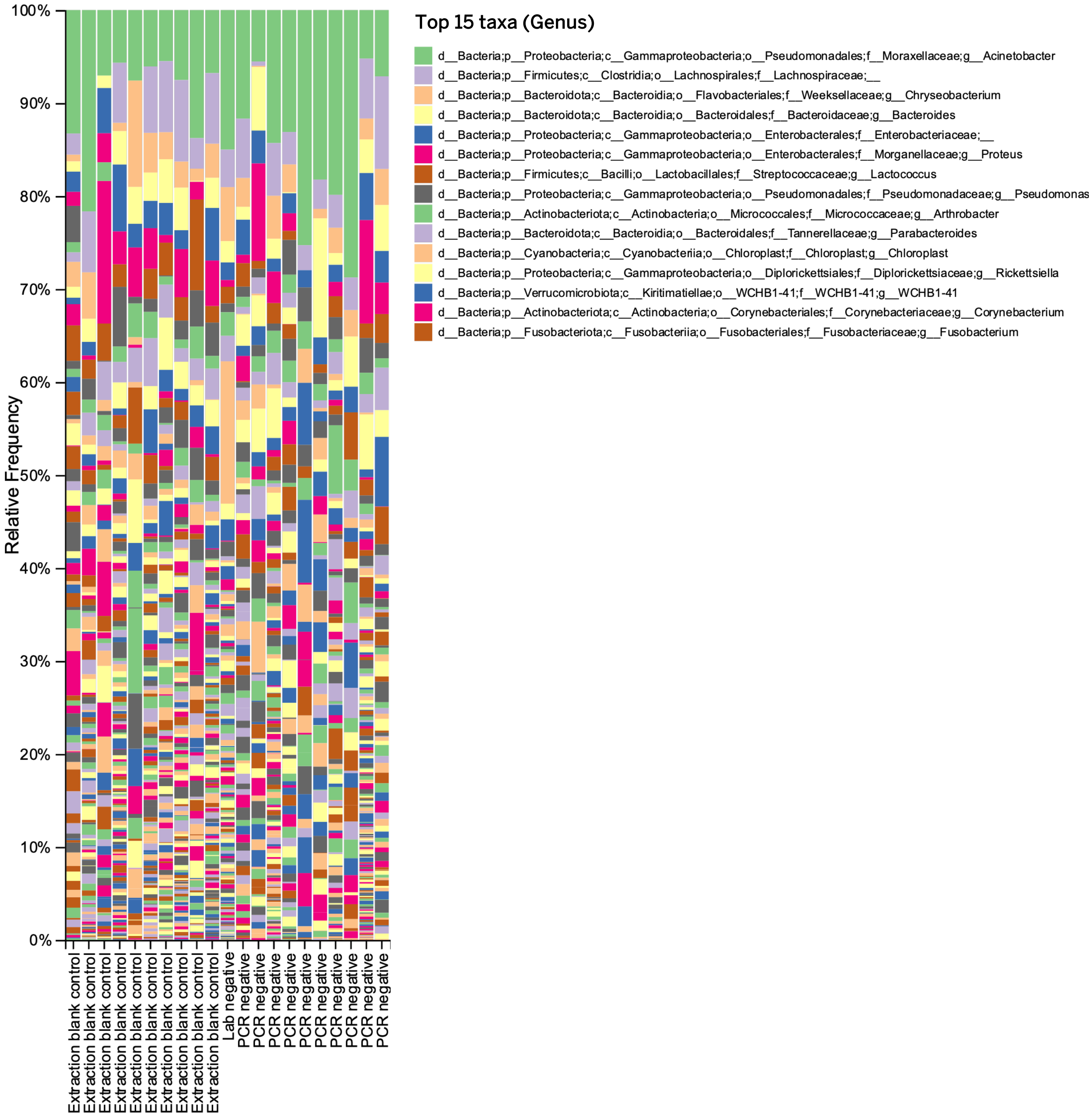


**Supplementary figure S9: Taxonomic composition of negative controls prior to contaminant removal.** Taxa are displayed to the genus level. Top 15 most prevalent taxa are listed to the right of the plot.
